# Evolution of the proinflammatory receptor TREM-1 in mammals reveals signatures of pathogen-driven conflict

**DOI:** 10.64898/2026.08.05.743136

**Authors:** Omoshola Aleru, Kaitlyn E. Bunn, Matthew F. Barber

## Abstract

Animal immune cells express a range of surface receptors that promote the detection of diverse host and pathogen-derived molecules. The triggering receptors expressed on myeloid cells (TREMs) encompass a family of cell surface receptors involved in the modulation of immune signaling cascades. Mammalian TREM-1 has emerged as a critical mediator of antibacterial immune defense and inflammatory disease, yet much remains unknown regarding its evolution and relevant molecular interactions. Here we applied a comparative phylogenetic approach to investigate patterns of divergence and natural selection among mammalian TREM-1 orthologs. We identify evidence of repeated positive selection acting within the extracellular ligand binding domain of TREM-1 among primates, rodents, and particularly bats. Structural simulations further suggest that genetic variation in TREM-1 impacts recognition of putative host and microbial ligands, with implications for downstream signaling functions. Together our findings identify patterns of rapid divergence in mammalian TREM-1, suggesting a history of evolutionary conflict in response to pathogen antagonism.

## INTRODUCTION

Cell surface receptors play key roles in host immune defense through detection of both self (endogenous) and non-self (exogenous) ligands (Takeuchi & Akira 2010; Arpaia & Barton 2013; Bloes et al. 2015; Weiß & Kretschmer 2018). The activation of immune cell surface receptors can lead to several downstream responses such as inflammatory cytokine production, leukocyte recruitment, release of antimicrobial compounds, tissue repair, or initiation of adaptive immune responses (Takeuchi & Akira 2010). Pattern recognition receptors (PRRs), for example, encompass a wide range of distinct receptor families which have evolved to detect microbial associated molecular patterns, or MAMPs. Conversely, some immune receptors are tuned to detect host-derived damage-associated molecular patterns (DAMPs), which are typically associated with tissue injury or host cell death. The triggering receptor expressed on myeloid cells (TREM) gene family comprises a group of cell surface receptors that play key roles in the modulation of vertebrate immune signaling pathways (Klesney-Tait et al. 2006; Colonna 2023).

TREM family members are differentially expressed on leukocytes (white blood cells) of the myeloid lineage, including neutrophils, macrophages, monocytes, and dendritic cells (Colonna 2023). Activation of TREMs by both exogenous and endogenous ligands stimulates a range of outcomes including cell migration, phagocytosis, and cytokine production (Klesney-Tait et al. 2006; Colonna 2023). Distinct TREM family members can promote both pro-inflammatory (activating) and anti-inflammatory (regulatory) signaling cascades depending on their binding interactions. All TREM proteins contain a single N-terminal extracellular V-type immunoglobulin-like (IgV-like) domain, a transmembrane domain, and a short cytoplasmic domain. While most TREMs lack an obvious cytoplasmic signaling motif, the transmembrane domains of activating TREMs are capable of forming complexes with DNAX activating protein 12 (DAP-12) which contains an immunoreceptor tyrosine-based activation motif (ITAM) that promotes pro-inflammatory signaling (Fig. 1A). DAP-12 serves as a signal transducer for a range of receptors expressed by natural killer and myeloid cells and plays broad roles in microbial pathogen recognition, cell differentiation and immune signaling (Hamerman et al. 2009; Turnbull & Colonna 2007). DAP-12 associated TREMs, including the founding member TREM-1, are potent activators of inflammatory signaling and amplify signaling through other PRRs including Toll-like receptors (TLRs) (Arts et al. 2013; Haselmayer et al. 2009).

**Figure 1.**
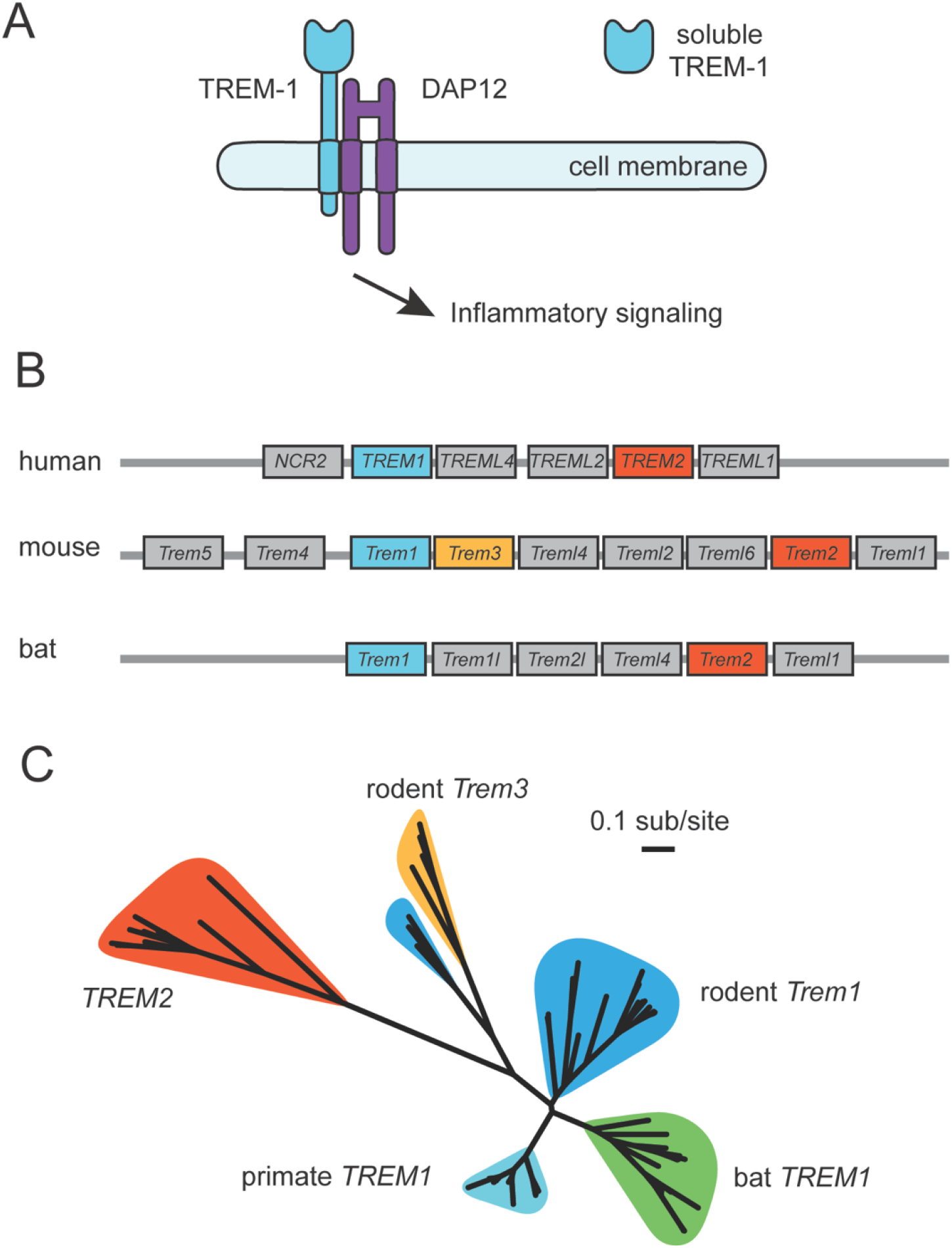
Divergence of TREM-1 in primates, rodents, and bats. (A) Schematic depicting TREM-1 (green) bound to co-receptor DAP-12 (purple), which is required for inflammatory signal activation. TREM-1 is known to be produced as both a cell surface protein (left) and a soluble extracellular protein (right). (B) Depiction of the TREM gene locus in representative human (*H. sapiens*), rodent (*M. musculus*), and bat (*M. lucifigus*) genomes. *TREM1* is denoted in green, *TREM2* in red, and *TREM3* in orange. (C) Maximum-likelihood unrooted phylogenetic tree of representative TREM-1, TREM-2, and TREM-3 orthologs from primates, rodents, and bats. Major clades are colored as follows: primate TREM-1 (cyan), rodent TREM-1 (blue), bat TREM-1 (green), TREM-2 (red), and TREM-3 (orange).

TREM-1 is expressed on neutrophils, monocytes, and certain macrophage subtypes (Arts et al. 2013; Colonna 2023). Functional and genetic studies have established TREM-1 as an important mediator of pathogen defense and sterile immunity. For example, TREM-1 signaling is highly upregulated during bacterial infections and in some cases contributes to pathogen clearance (Lin et al. 2014; Yang et al. 2015; Matos et al. 2021). TREM-1 activation also plays roles in non-infectious inflammatory disease states such as rheumatoid arthritis, psoriasis, and inflammatory bowel disease (Colonna 2023; Al Sawy et al. 2026; Schenk et al. 2007; Inanc et al. 2021; Hyder et al. 2013).

Polymorphisms in TREM-1 have also been implicated in susceptibility of humans to inflammatory diseases (Rivera-Chavez et al. 2006; de Jesus et al. 2023), yet the mechanisms underlying these observations remain unclear. These findings suggest that TREM-1 activity must be finely tuned to promote proper immune function while avoiding deleterious autoimmune conditions. These factors also make TREM-1 an attractive target for therapeutic intervention in both infectious and inflammatory disease states.

While past work has established important roles of TREM-1 in modulating host inflammatory signaling, many fundamental questions remain regarding its biology. For example, *bona fide* activating ligands of TREM-1 remain in question. The most promising candidate to date is peptidoglycan recognition protein 1 (PGLYRP1), a soluble protein released by activated neutrophils during degranulation (Read et al. 2015; Rickert et al. 2025). PGLYRP1 itself contributes to immune defense by binding bacterial-derived peptidoglycan, which can both interfer with pathogen replication as well as sequester this potent MAMP from other inflammatory PRRs (Gupta 2008).

These observations suggest that TREM-1 may sense active infection through the detection of host ligands like PGLYRP1. While TREM-1 signaling is upregulated during bacterial infections, it remains unclear whether any bacterial-derived MAMPs are capable of directly activating this receptor. In addition, some studies suggest that alternative splicing or proteolytic cleavage may be important regulators of TREM-1 function (Cao et al. 2017). TREM-1 has been found to be expressed as both a membrane-bound and a soluble extracellular protein (Allaouchiche & Boselli 2004; Molad & Lagovsky 2025). While soluble TREM-1 has been investigated as a useful biomarker of inflammatory state (Cao et al. 2017), the functional relevance of this secreted protein form remains unclear. Thus, while many studies have now implicated TREM-1 as an important inflammatory signal amplifier in health and disease, details regarding its precise functions remain enigmatic. Applying orthologonal research strategies to investigate the genetic diversity of TREM-1 could therefore yield useful new information for the field.

In this study we took a molecular phylogenetics approach to investigate patterns of divergence and natural selection in TREM-1 among primates, rodents, and bats. We identify evidence of repeated positive selection predominantly within the extracellular IgV-like domain of TREM-1, as well as strong conservation of the DAP12-interacting transmembrane region. Structural simulations of the TREM-1-PGLYRP1 interaction site suggest that rapid evolution of TREM-1 in distinct mammal lineages has likely had a major impact on ligand binding, as well as suggesting a history of adaptation driven by as-yet unknown pathogen binding interactions. Collectively this study provides an improved understanding of TREM gene family evolution and informs future studies to identify and characterize TREM-1 ligand interactions.

## RESULTS

### Phylogenetic relatedness of mammalian TREMs

To begin to investigate the evolution of TREM-1 in mammals, we assembled a series of TREM family homologs from representative primates, rodents, and bats. We selected these three taxa as they are both species-rich and important sources of zoonotic pathogens. The locus encoding TREM genes is well-conserved among these three groups, containing orthologs of TREM-1, TREM-2, and a series of TREM-like (TREML) homologs (Fig. 1B). Rodents also possess a unique paralog, TREM-3. We inferred a maximum-likelihood phylogenetic tree of representative primate, rodent, and bat paralogs focusing on TREM-1, TREM-2, and (for rodents) TREM3 (Fig. 1C). The phylogeny was generally concordant with known gene and species relationships in this group, with the exception that a subset of annotated rodent TREM-1 orthologs were inferred to form a clade with TREM3 orthologs (Fig. 1C). This observation could reflect mis-annotation of these rodent TREM-1 orthologs, or alternatively be explained by gene conversion between these loci in rodents giving rise to the inferred shared ancestry.

Given this phylogenetic discordance, this subset of rodent TREM-1 orthologs were omitted from subsequent phylogenetic analyses.

### Signatures of recurrent positive selection within the TREM-1 IgV-like domain

Many immune receptors, including those expressed on myeloid cells, have been found to undergo repeated positive selection in diverse animal lineages (Radwan et al. 2020; Daugherty & Malik 2012; Demogines et al. 2013; Paterson et al. 2021; Carey et al. 2021; Paterson et al. 2023). Evidence suggests that selection may be due to a range of pressures, such as adaptation to recognize evolving pathogen ligands, or evasion of pathogen receptors and virulence factors that target these proteins to promote infection or immune evasion (Radwan et al. 2020; Paterson et al. 2021; Boguslawski et al. 2020). To assess whether TREM-1 orthologs from primates, rodents, or bats exhibit signatures of repeated positive selection, we applied a series of phylogenetic analyses from the HyPhy software package (Pond et al. 2005; Weaver et al. 2018). We particularly focused on approaches designed to detect signatures of positive selection at single sites within protein coding genes, inferred from the rate of nonsynonymous to synonymous substitutions (dN/dS). Our dataset included TREM-1 orthologs from 23 primates, 16 rodents, and 18 bat species. We used three independent statistically analyses to assess evidence of selection: MEME, FUBAR, and FEL. We chose to further investigate all sites that were identified by two or more of these analyses (Fig. 2A). Notably, we detected sites with elevated dN/dS in all three mammalian lineages. These observations suggest that TREM-1 has indeed been subject to repeated positive selection in multiple mammalian lineages over millions of years. Looking more closely at domains where these rapidly evolving sites lie, we noted that the extracellular IgV-like domain appeared to be a “hotspot” of positive selection in both primates and bats (Fig. 2A). Four of six rapidly evolving sites in primates were found in the IgV-like domain, as well as 14 out of 15 sites in bats. While six sites were identified with elevated dN/dS in our rodent dataset, only two of these map to the IgV-like domain. Comparing between clades, we noted overlap in rapidly evolving sites. For example, three sites under positive selection in rodents align to the same position in bat TREM-1. Similarly, one rapidly evolving site identified in primates also overlaps with a rapidly evolving position in bats. To gain a structural perspective on these observations, we mapped rapidly evolving sites from all three clades onto predicted IgV-like domain structures from humans, mice, and bats (Fig. 2B). For humans we leveraged a previously solved crystal structure of the TREM-1 IgV-like domain, while mouse and bat TREM-1 IgV-like structures were predicted using AlphaFold3 (Abramson et al. 2024). We noted that sites subject to positive selection map to different surfaces along the IgV-like domain (Fig.2B). This was particularly true of bat TREM-1, where numerous sites mapped along nearly every surface of the domain. Together these results suggest that the TREM-1 gene has experienced repeated positive selection in several mammalian lineages, with particularly strong support within the IgV-like domain of bats.

**Figure 2.**
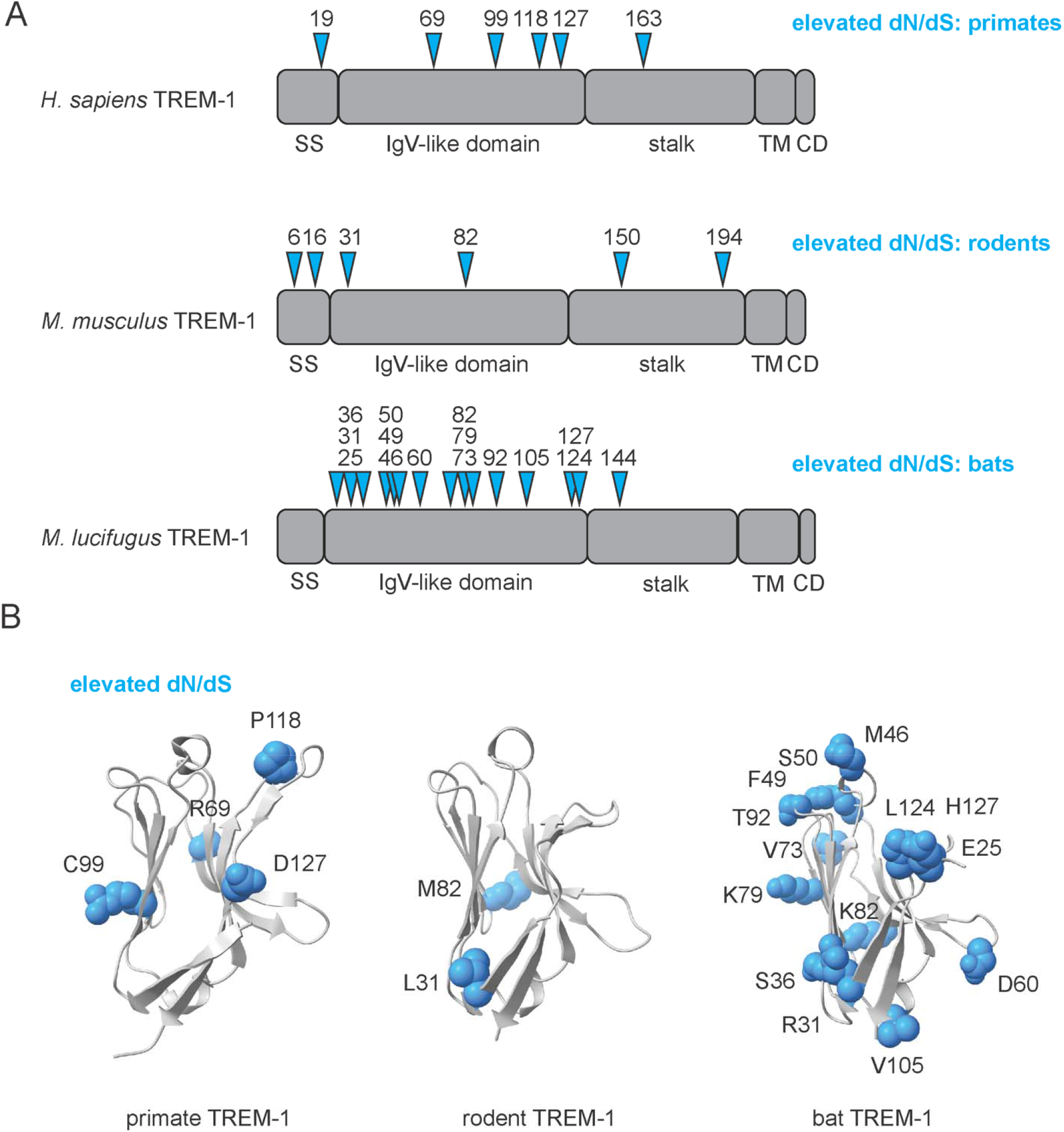
Evidence of repeated positive selection in mammalian TREM-1. (A) Diagrams of the TREM-1 protein from primates, rodents, and bats. Blue arrows identify sites exhibiting signatures of positive selection in at least two independent tests using the HyPhy software package. SS: signal sequence; TM: trans-membrane domain; CD: cytoplasmic domain. (B) Structures of the IgV-like domain from three representative mammals with side-chains of amino acid positions exhibiting signatures of positive selection mapped for primates, rodents, and bats (blue). *H. sapiens* TREM-1 crystal structure is shown as primate representative (PDB: 1Q8M). The TREM-1 IgV-like domains of *M. musculus* and *M. lucifigus* were predicted using AlphaFold as representatives for rodents and bats, respectively.

### Conservation of the DAP12-interacting site among TREM-1 orthologs

TREM proteins have been shown to possess an unusual mode of activation by the co-receptor DAP-12. TREM-DAP-12 physical interaction is mediated by the transmembrane domains of the proteins, while DAP-12’s cytoplasmic ITAM mediates inflammatory signaling downstream of TREM ligand binding. Specifically, a single lysine residue in the TREM-1 transmembrane domain has been shown to promote electrostatic interactions with an aspartic acid residue in the DAP12 transmembrane domain (Bouchon et al. 2000). Given observations of rapid amino acid divergence and selection in the TREM-1 IgV-like domain, we investigated whether the TREM-1 transmembrane domain exhibited any evidence of diversification and possible loss of DAP12-mediated activation. Alignment of the TREM-1 transmembrane domain from a subset of primates, rodents, and bats revealed limited sequence conservation in this region, likely reflecting relaxed constraint on this region (Fig. 3A). The transmembrane domain was enriched for hydrophobic amino acids - particularly valine, lysine, isoleucine, and phenylalanine - consistent with such regions that mediate interactions with hydrophobic lipid membranes. Despite the low level of general sequence conservation, we noted that the lysine residue which mediates DAP12 interactions was conserved across the surveyed species (Fig. 3A). These findings are consistent with long-term maintenance of the DAP12-interacting region in mammalian TREM-1 that mediates inflammatory signal transduction.

**Figure 3.**
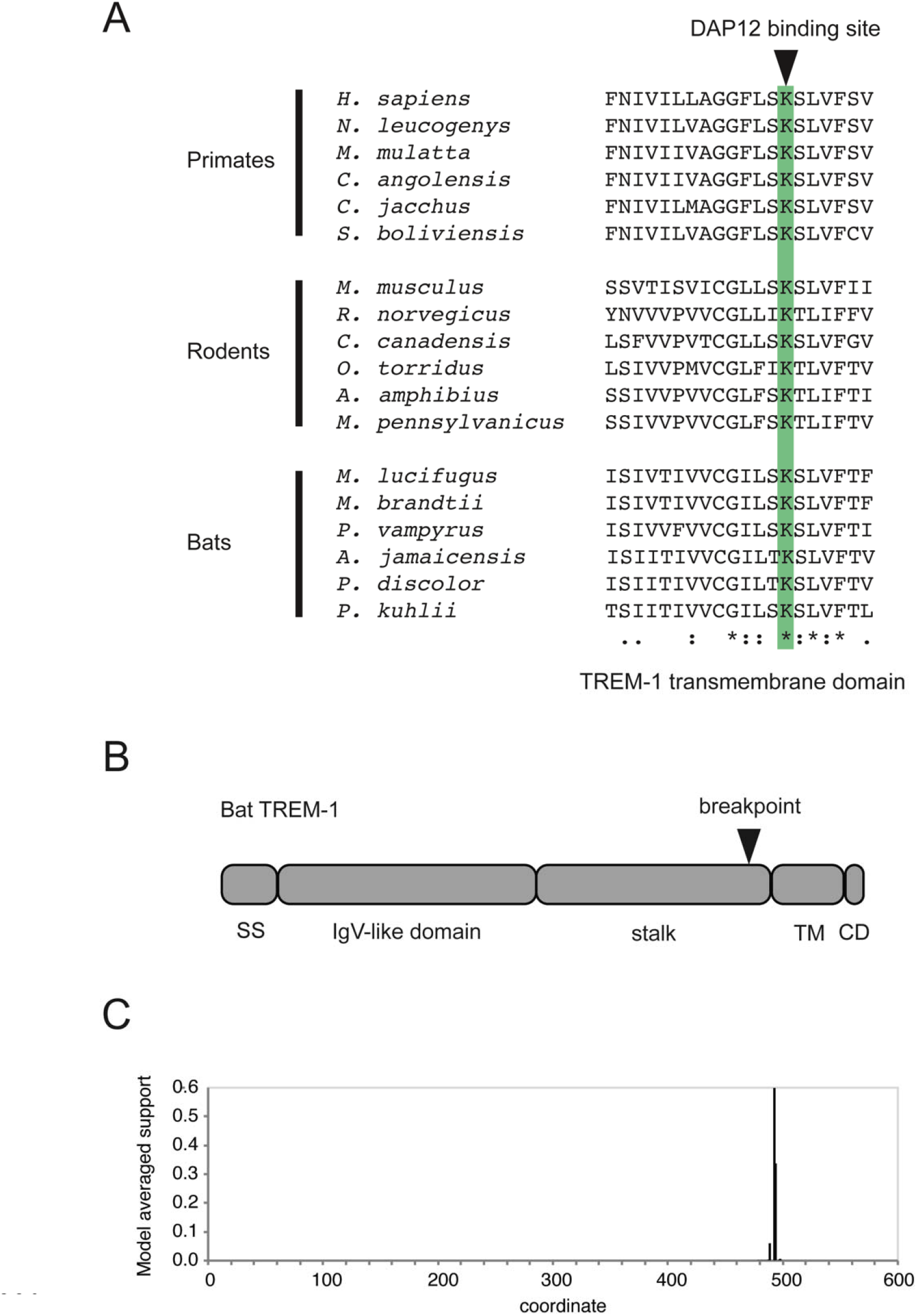
Patterns of conservation and recombination in mammalian TREM-1. (A) Amino acid sequence alignment of the transmembrane domains of representative mammalian TREM-1 orthologs. The position of the lysine residue which mediates interactions with DAP-12 is highlighted in green. (B) Diagram of the bat TREM-1 protein with position of the predicted recombination breakpoint. (C) Model average support as determined using the GARD algorithm to identify potential recombination breakpoints in bat TREM-1, as shown in (B).

### Evidence of recombination in bat TREM-1

Previous work by numerous groups has found that immune receptors, particularly those possessing one or more immunoglobulin domains, undergo recombination and gene conversion of homologous extracellular ligand-binding domains (Kawamura et al. 1992; Baker et al. 2022). In addition to amino acid substitutions inferred by dN/dS, recombination is an alternative evolutionary process that can modify receptor binding functions (Daugherty & Zanders 2019). To investigate the potential for recombination acting within mammalian TREM-1, we applied the GARD algorithm from the HyPhy software package (Kosakovsky Pond et al. 2006). GARD functions by searching for positions of discordance in phylogenies generated from segments of a gene in order to identify possible recombination breakpoints. Applying this approach to our TREM-1 datasets, we identified strong evidence for a single breakpoint in bats, but not primates or rodents (Fig. 3B, C). This breakpoint was located in the C-terminal portion of the TREM-1 stalk domain, near the transmembrane domain. These findings are consistent with recombination functioning to swap or modify the extracellular region of TREM-1, which could alter ligand binding function while maintaining the conserved DAP12 binding activity.

### Modeling potential TREM-1-ligand interactions

Evidence of repeated positive selection in TREM-1 raises the question of what might be driving such drastic variation. We considered two possibilities. The first is that selection could occur in response to changes in one or more TREM-1 activating ligands, which themselves may be host or pathogen-derived. The second possibility is that TREM-1 variation could be driven by evasion of pathogen antagonists that bind to TREM-1 in order to promote infection or pathogenesis. These could include viral entry receptors, bacterial toxins, or other virulence factors. However, little is known regarding any specific molecular interactions involving TREM-1, with host or pathogen. A current promising lead is peptidoglycan recognition protein 1 (PGLYRP1), a protein released by neutrophil granules in response to infection. PGLYRP1 is believed to function by binding to bacterial-derived cell wall peptidoglycan molecules, coating the bacterial cell surface and triggering cell death (Gupta 2008). It is also possible PGLYRP1 could aid in preventing deleterious immune activation during infection by sequestering peptidoglycan from other PRRs. Previous studies have shown that PGLYRP1 is capable of binding to TREM-1 and promoting signal transduction (Read et al. 2015).

However, the molecular basis of this interaction remains unclear.

To gain some insight into the possible consequences of TREM-1 evolution for ligand binding, we used AlphaFold to predict the structure of the protein-protein interaction between PGLYRP1 and the TREM-1 IgV-like domain. We then mapped sites with elevated dN/dS onto this predicted protein complex (Fig. 4A). We observed that numerous rapidly evolving sites in TREM-1 lay at the predicted interface with PGLYRP1, suggesting that TREM-1 evolution could indeed have consequences for host ligand interactions. Collectively these results support the hypothesis that repeated positive selection of TREM-1 impacts recognition of the host-derived activating ligand PGLYRP1.

**Figure 4.**
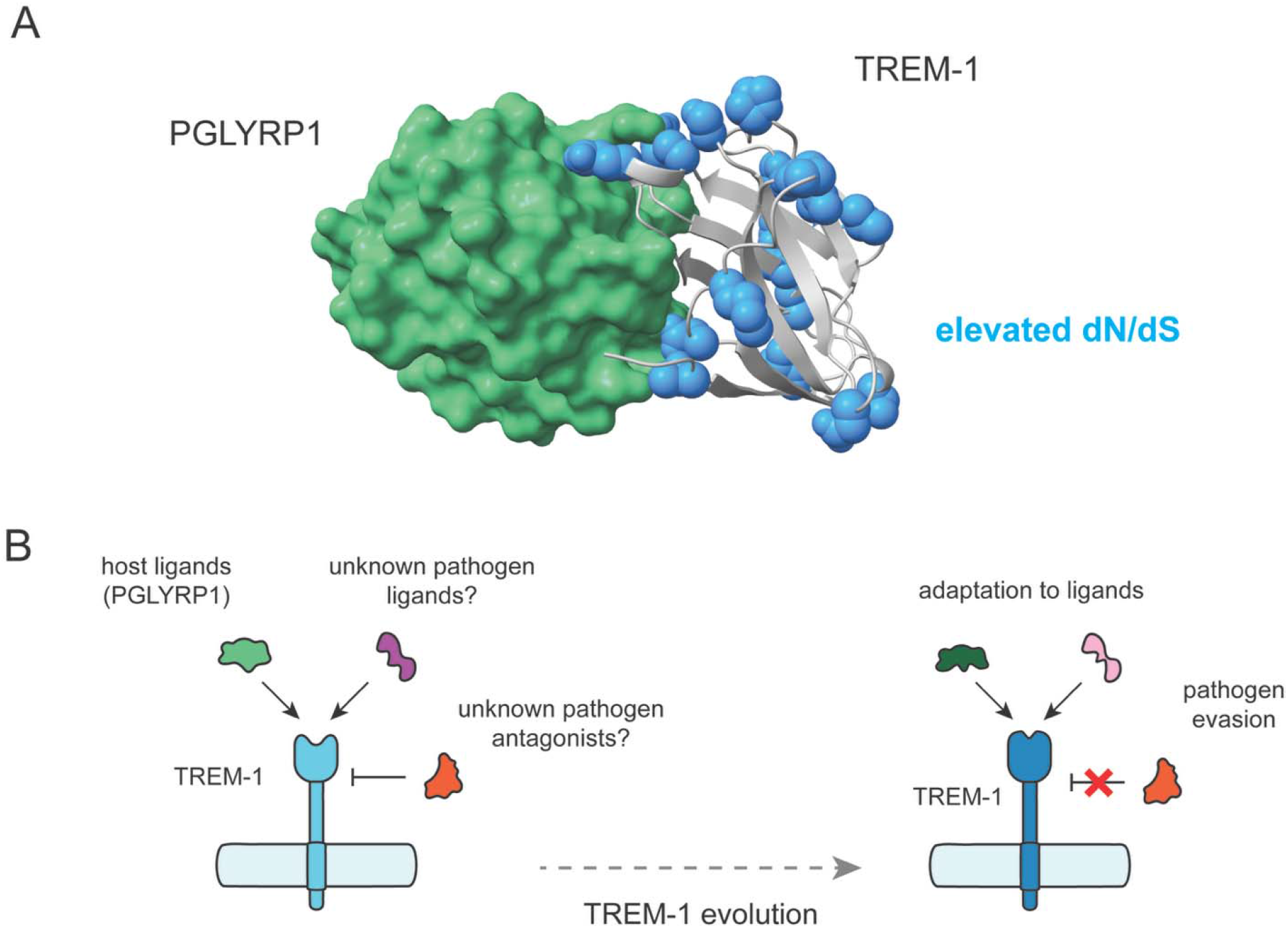
Potential consequences of rapid evolutionary divergence in mammalian TREM-1. (A) Predicted interaction between human TREM-1 (grey) and PGLYRP-1 (green) was determined using AlphaFold. Sites in TREM-1 exhibiting signatures of positive selection among primates, rodents, or bats are highlighted in blue. (B) Schematic depicting potential molecular interactions shaping TREM-1 evolution.

## DISCUSSION

Cell surface immune receptors have been found to be engaged in host-pathogen evolutionary conflicts across diverse taxa (Daugherty & Malik 2012; Sironi et al. 2015; Barber & Fitzgerald 2024). Natural selection in such proteins can serve to promote targeting of rapidly evolving pathogens (Tenthorey et al. 2017; Adrian et al. 2019; Radwan et al. 2020; Kohler et al. 2020; Paterson et al. 2021). Conversely, such variation may also mediate evasion of pathogen receptors or virulence factors (Boguslawski et al. 2020; Baker et al. 2022). While genetic and functional evidence points to important functions of TREMs in host immune defense and inflammation, many fundamental questions remain outstanding. In this study we find that TREM-1 orthologs across multiple mammalian lineages exhibit signatures of repeated positive selection, with rapid amino acid variation identified in the extracellular IgV-like domain.

Extracellular IgV-like domains in many immune receptors mediate direct binding of both exogenous and endogenous ligands. However, the *bona fide* binding partners of TREM-1 have remained elusive. This raises the question of what may be driving repeated variation in the TREM-1 IgV-like domain. Several studies have now provided support that PGLYRP-1, a protein released by activated neutrophils, is an activating ligand for TREM-1 (Read et al. 2015; Rickert et al. 2025). Our structural modeling predictions suggest that rapid variation of the IgV-like domain would likely impact interactions with PGLYRP-1, particularly in bats (Fig. 4A). Whether this domain is evolving in response to PGLYRP-1 variation, or conversely whether PGLYRP-1 may co-evolve with TREM-1 to maintain this interaction in mammals remains unknown. Future studies that identify new ligands for TREM-1 or further clarify the molecular basis for PGLYRP-1-mediated activation of TREM-1 would aid in resolving these questions.

Conservation of the TREM-1 transmembrane lysine residue suggests that purifying selection has acted to maintain interactions with DAP12 in this otherwise highly variable domain (Fig. 3B, C). These findings indicate that TREM-1’s proinflammatory activity is conserved in diverse mammals. Evidence of a recombination breakpoint in bat TREM-1 suggests, however, that the evolution of TREM family members may be actively shaped by processes such as gene conversion. Our observation of phylogenetic discordance among a subset of rodent TREM-1 orthologs (Fig. 1C) could be indicative of such recombination events. Previous work by our group and others has shown how repeated gene conversion in cell surface or cytoplasmic immune receptors can serve to modulate microbial binding interactions, both positively or negatively, to promote host defense (Tenthorey et al. 2014; Daugherty et al. 2016; Mitchell et al. 2015; Baker et al. 2022).

Whether recombination might play a role in modifying TREM IgV-like domain function or perhaps swapping signal activation domains remains to be determined.

In conclusion, our study provides new genetic and evolutionary information regarding patterns of divergence in the mammalian inflammatory signal amplifier TREM-1. Future studies aimed at discerning the functional impacts of TREM substitutions or recombination will ultimately provide a more complete view of potential drivers of such variation. Moreover, the lack of knowledge regarding physiologically relevant ligands for TREM-1 may be informed by incorporating knowledge regarding the diversification of this gene across mammals. Together this work advances our understanding regarding patterns of evolution among key receptors of the animal innate immune system.

## METHODS

### Comparative genetics and tree building

Orthologs of TREM-1, TREM-2, and TREM-3 were identified via searches of the NCBI Gene database as well as BLAST searches of primate, rodent, and bat reference genomes via the NCBI BLAST webserver. Full length coding sequences for annotated human or mouse TREM orthologs were used as query sequences. A full list of orthologs identified are provided in Supplemental Data. Phylogenetic trees were inferred using mammalian TREM ortholog coding sequences as described above using the PhyML 3.0 web browser (http://www.atgc-montpellier.fr/phyml/) with default settings (Guindon et al. 2010).

### Phylogenetic analyses

For the following methods, each mammalian group of TREM1 coding sequences (primates, rodents, and bats) were analyzed separately. Multiple sequence alignments of the full-length coding sequence were generated using MUSCLE (Edgar 2004).

Alignment gaps were removed to maintain reading frame and codon structure. Sites exhibiting signatures of positive selection were identified using the HyPhy package programs MEME, FUBAR, and FEL (Kosakovsky Pond & Frost 2005; Murrell et al. 2012, 2013) as implemented on the Datamonkey 2.0 webserver (Weaver et al. 2018). Sites highlighted in Figure 2 reflect positions that were identified in two or more of these analyses using default settings. The HyPhy GARD analysis was applied to detect evidence of recombination breakpoints for all three groups (Kosakovsky Pond et al. 2006).

### Structural modeling

Protein structures were visualized using ChimeraX-1.9 (Pettersen et al. 2021). Human TREM-1 was based on the published crystal structure (Radaev et al. 2003), whereas mouse and bat TREM-1 IgV-like domain structures were generated using AlpahFold3 (Abramson et al. 2024) implemented through the AlphaFold server (https://alphafoldserver.com/). The predicted protein interaction between human TREM-1 and human PGLYRP-1 was also generated using AlphaFold3. Raw data associated with predicted structures are provided in Supplemental Data.

## ACKNOWLEDGMENTS

We thank Michael Harms and members of the Barber lab for helpful discussions. This work was supported by National Institutes of Health grant R35GM133652 and R35GM158176 (to M.F.B.). O.A. is a recipient of an NIH Molecular Biology and Biophysics training grant (T32GM007759).

## COMPETING INTERESTS STATEMENT

The authors declare no competing financial interests.

## Notes

### Competing Interest Statement

The authors have declared no competing interest.

### Summary of Updates

Addition of supplemental data files used for phylogenetic analysis of primate, rodent, and bat TREM-1 orthologs.

## REFERENCES

Abramson J et al. 2024. Accurate structure prediction of biomolecular interactions with AlphaFold 3. Nature. 630:493–500.

Adrian J, Bonsignore P, Hammer S, Frickey T, Hauck CR. 2019. Adaptation to Host-Specific Bacterial Pathogens Drives Rapid Evolution of a Human Innate Immune Receptor. Current Biology. 29:616–630.e5. doi: 10.1016/j.cub.2019.01.058.

Al Sawy ER, Saber MM, Nassar NN, El Sayed NS. 2026. TREM-1 receptor: A key player in inflammatory diseases. Curr. Mol. Pharmacol. 19:41–47.

Allaouchiche B, Boselli E. 2004. Soluble TREM-1 and the diagnosis of pneumonia. N. Engl. J. Med. 350:1904–5; author reply 1904-5.

Arpaia N, Barton GM. 2013. The impact of Toll-like receptors on bacterial virulence strategies. Curr. Opin. Microbiol. 16:17–22.

Arts RJW, Joosten LAB, van der Meer JWM, Netea MG. 2013. TREM-1: intracellular signaling pathways and interaction with pattern recognition receptors. J. Leukoc. Biol. 93:209–215.

Baker EP et al. 2022. Evolution of host-microbe cell adherence by receptor domain shuffling. Elife. 11:e73330.

Barber MF, Fitzgerald JR. 2024. Mechanisms of host adaptation by bacterial pathogens. FEMS Microbiol. Rev. 48. doi: 10.1093/femsre/fuae019.

Bloes DA, Kretschmer D, Peschel A. 2015. Enemy attraction: bacterial agonists for leukocyte chemotaxis receptors. Nat. Rev. Microbiol. 13:95–104.

Boguslawski KM et al. 2020. Exploiting species specificity to understand the tropism of a human-specific toxin. Sci Adv. 6:eaax7515.

Bouchon A, Dietrich J, Colonna M. 2000. Cutting edge: inflammatory responses can be triggered by TREM-1, a novel receptor expressed on neutrophils and monocytes. J. Immunol. 164:4991–4995.

Cao C, Gu J, Zhang J. 2017. Soluble triggering receptor expressed on myeloid cell-1 (sTREM-1): a potential biomarker for the diagnosis of infectious diseases. Front. Med. 11:169–177.

Carey CM, Apple SE, Hilbert ZA, Kay MS, Elde NC. 2021. Diarrheal pathogens trigger rapid evolution of the guanylate cyclase-C signaling axis in bats. Cell Host Microbe. 29:1342–1350.e5.

Colonna M. 2023. The biology of TREM receptors. Nat. Rev. Immunol. 23:580–594.

Daugherty MD, Malik HS. 2012. Rules of engagement: molecular insights from host-virus arms races. Annu. Rev. Genet. 46:677–700.

Daugherty MD, Schaller AM, Geballe AP, Malik HS. 2016. Evolution-guided functional analyses reveal diverse antiviral specificities encoded by IFIT1 genes in mammals. Elife. https://elifesciences.org/content/5/e14228v1.

Daugherty MD, Zanders SE. 2019. Gene conversion generates evolutionary novelty that fuels genetic conflicts. Curr. Opin. Genet. Dev. 58–59:49–54.

Demogines A, Abraham J, Choe H, Farzan M, Sawyer SL. 2013. Dual host-virus arms races shape an essential housekeeping protein. PLoS Biol. 11:e1001571.

Edgar RC. 2004. MUSCLE: a multiple sequence alignment method with reduced time and space complexity. BMC Bioinformatics. 5:113.

Guindon S et al. 2010. New algorithms and methods to estimate maximum-likelihood phylogenies: assessing the performance of PhyML 3.0. Syst. Biol. 59:307–321.

Gupta D. 2008. Peptidoglycan Recognition Proteins?Maintaining Immune Homeostasis and Normal Development. Cell Host Microbe. 3:273–274.

Hamerman JA, Ni M, Killebrew JR, Chu CL, Lowell CA. 2009. The expanding roles of ITAM adapters FcRγ and DAP12 in myeloid cells. Immunological Reviews. 232:42–58.

Haselmayer P et al. 2009. Signaling pathways of the TREM-1- and TLR4-mediated neutrophil oxidative burst. J. Innate Immun. 1:582–591.

Hyder LA et al. 2013. TREM-1 as a potential therapeutic target in psoriasis. J. Invest. Dermatol. 133:1742–1751.

Inanc N et al. 2021. Elevated serum TREM-1 is associated with periodontitis and disease activity in rheumatoid arthritis. Sci. Rep. 11:2888.

de Jesus MCS et al. 2023. Influence of trem-1 gene polymorphisms on cytokine levels during malaria by Plasmodium vivax in a frontier area of the Brazilian Amazon. Cytokine. 169:156264.

Kawamura S, Saitou N, Ueda S. 1992. Concerted evolution of the primate immunoglobulin alpha-gene through gene conversion. J. Biol. Chem. 267:7359–7367.

Klesney-Tait J, Turnbull IR, Colonna M. 2006. The TREM receptor family and signal integration. Nature Immunology. 7:1266–1273.

Kohler KM et al. 2020. A Rapidly Evolving Polybasic Motif Modulates Bacterial Detection by Guanylate Binding Proteins. MBio. 11. doi: 10.1128/mBio.00340-20.

Kosakovsky Pond SL, Frost SDW. 2005. Not so different after all: a comparison of methods for detecting amino acid sites under selection. Mol. Biol. Evol. 22:1208–1222.

Kosakovsky Pond SL, Posada D, Gravenor MB, Woelk CH, Frost SDW. 2006. Automated phylogenetic detection of recombination using a genetic algorithm. Mol. Biol. Evol. 23:1891–1901.

Lin Y-T et al. 2014. TREM-1 promotes survival during Klebsiella pneumoniae liver abscess in mice. Infect. Immun. 82:1335–1342.

Matos A de O, Dantas PHDS, Silva-Sales M, Sales-Campos H. 2021. TREM-1 isoforms in bacterial infections: to immune modulation and beyond. Crit. Rev. Microbiol. 47:290– 306.

Mitchell PS, Young JM, Emerman M, Malik HS. 2015. Evolutionary Analyses Suggest a Function of MxB Immunity Proteins Beyond Lentivirus Restriction. PLoS Pathog. 11:e1005304–21.

Molad Y, Lagovsky I. 2025. Soluble TREM-1 ameliorates gouty arthritis by selective inhibition of proinflammatory cytokines and chemokines without affecting TGFβ production. Clin. Exp. Rheumatol. 43:1252–1258.

Murrell B et al. 2012. Detecting Individual Sites Subject to Episodic Diversifying Selection. PLoS Genet. 8:e1002764.

Murrell B et al. 2013. FUBAR: a fast, unconstrained bayesian approximation for inferring selection. Mol. Biol. Evol. 30:1196–1205.

Paterson NM et al. 2023. Dynamic Evolution of Bacterial Ligand Recognition by Formyl Peptide Receptors. Genome Biol. Evol. 15. doi: 10.1093/gbe/evad175.

Paterson NM, Al-Zubieri H, Barber MF. 2021. Diversification of CD1 Molecules Shapes Lipid Antigen Selectivity. Mol. Biol. Evol. 38:2273–2284.

Pettersen EF et al. 2021. UCSF ChimeraX: Structure visualization for researchers, educators, and developers. Protein Sci. 30:70–82.

Pond SLK, Frost SDW, Muse SV. 2005. HyPhy: hypothesis testing using phylogenies. Bioinformatics. 21:676–679.

Radaev S, Kattah M, Rostro B, Colonna M, Sun PD. 2003. Crystal structure of the human myeloid cell activating receptor TREM-1. Structure. 11:1527–1535.

Radwan J, Babik W, Kaufman J, Lenz TL, Winternitz J. 2020. Advances in the Evolutionary Understanding of MHC Polymorphism. Trends Genet. 36:298–311.

Read CB et al. 2015. Cutting Edge: identification of neutrophil PGLYRP1 as a ligand for TREM-1. J. Immunol. 194:1417–1421.

Rickert DM et al. 2025. A dual role for PGLYRP1 in host defense and immune regulation during B. pertussis infection. bioRxivorg. doi: 10.1101/2025.09.26.678899.

Rivera-Chavez FA, Horton JW, Minei JP. 2006. A Trem-1 polymorphism influences mortality and sepsis severity in burn patients. Shock. 25:92.

Schenk M, Bouchon A, Seibold F, Mueller C. 2007. TREM-1--expressing intestinal macrophages crucially amplify chronic inflammation in experimental colitis and inflammatory bowel diseases. J. Clin. Invest. 117:3097–3106.

Sironi M, Cagliani R, Forni D, Clerici M. 2015. Evolutionary insights into host-pathogen interactions from mammalian sequence data. Nat. Rev. Genet. 16:224–236.

Takeuchi O, Akira S. 2010. Pattern recognition receptors and inflammation. Cell. 140:805–820.

Tenthorey JL et al. 2017. The structural basis of flagellin detection by NAIP5: A strategy to limit pathogen immune evasion. Science. 358:888–893.

Tenthorey JL, Kofoed EM, Daugherty MD, Malik HS, Vance RE. 2014. Molecular basis for specific recognition of bacterial ligands by NAIP/NLRC4 inflammasomes. Mol. Cell. 54:17–29.

Turnbull IR, Colonna M. 2007. Activating and inhibitory functions of DAP12. Nature Reviews Immunology. 7:155–161.

Weaver S et al. 2018. Datamonkey 2.0: a modern web application for characterizing selective and other evolutionary processes. Mol. Biol. Evol. doi: 10.1093/molbev/msx335.

Weiß E, Kretschmer D. 2018. Formyl-Peptide Receptors in Infection, Inflammation, and Cancer. Trends Immunol. 39:815–829.

Yang C et al. 2015. TREM-1 signaling promotes host defense during the early stage of infection with highly pathogenic Streptococcus suis. Infect. Immun. 83:3293–3301.

